# Arbitrium systems control lysis/lysogeny through the regulation of small antirepressor proteins

**DOI:** 10.1101/2025.11.23.689978

**Authors:** Stav Kabel, Shira Omer Bendori, Tom Borenstein, Polina Guler, Francisca Gallego del Sol, Javier Mancheño-Bonilla, Alberto Marina, Avigdor Eldar

## Abstract

Many temperate *Bacillus* phages utilize the arbitrium signaling system to control lysis/lysogeny decisions. While the function of the arbitrium signal AimP and its receptor AimR are well known, it is unclear how they control lysis in most arbitrium systems. Here, we show that a large majority of arbitrium systems are embedded in an extended module with three additional components; A small antirepressor protein (AimX), the phage repressor (AimC) and an adjacent cro-like protein (AimL). AimR-dependent activation of AimX is necessary for lysis both during infection and lytic induction. Molecular analysis suggests that AimX directly binds AimC and prevents its oligomerization and binding to its regulated *aimL* promoter. Our work therefore uncovers the main mechanism by which arbitrium systems regulate lysis and point to the central role of small proteins in phage decision making.

**Graphical abstract:** 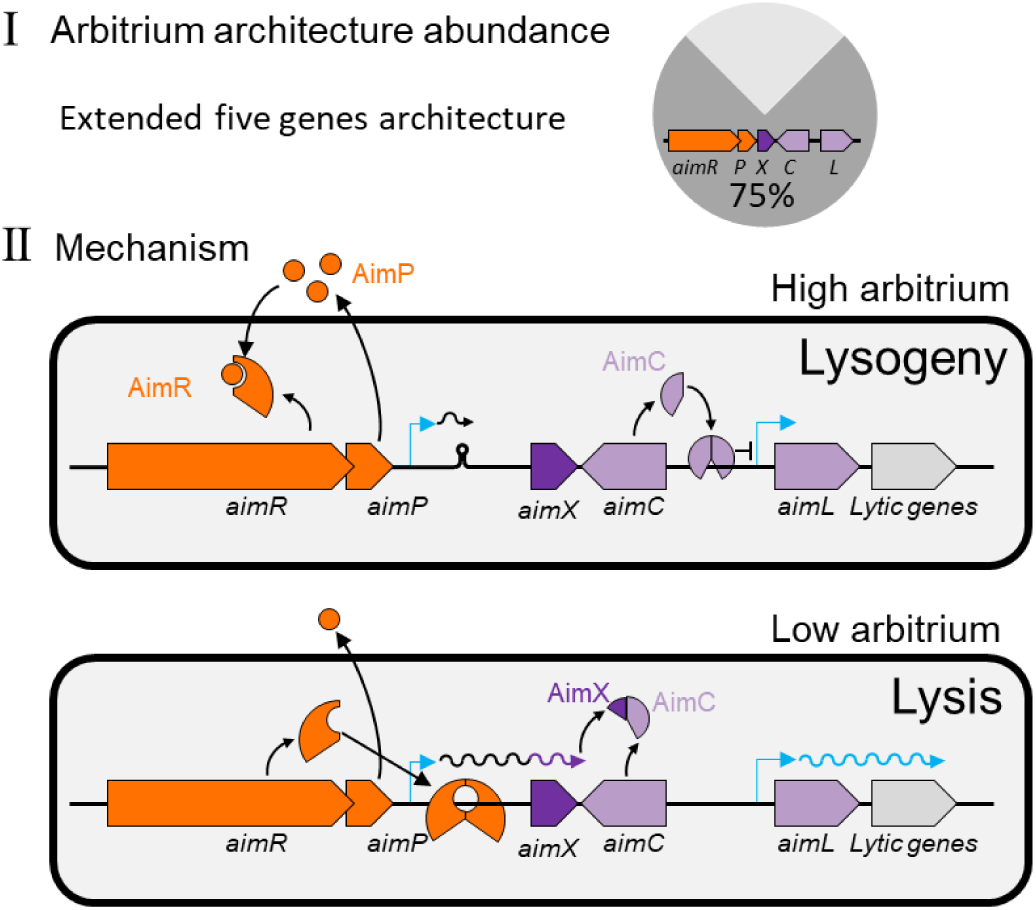

## Introduction

Temperate bacteriophages (or phages) carefully control the transition between their quiescent (lysogenic) and virulent (lytic) life-cycle stages^1–4^. This decision-making process relies on information of host conditions and on the surrounding population^5,6^. Upon infection, phages have been shown to incorporate information regarding phage density by sensing phage coinfection. Phages are also able to sense the physiological state of the host through diverse mechanisms and modify the outcome of infections accordingly ^2,6,7^.

The transition from lysogeny back to lysis, termed lytic induction, has similarly been shown to be influenced by both host-specific and population-wide factors^2,6–8^. Specifically, DNA damage is found to activate lysis in a majority of phages. In phage lambda, the phage lysogeny master repressor, CI, contain an auto-cleaving domain that responds to DNA-damage dependent activated RecA. CI degradation then allows the activation of the early lytic Cro transcriptional regulator and other co-cistronic lytic genes ^6^. In other phages, DNA damage dependent degradation of the bacterial LexA SOS regulator, leads to activation of phage antirepressors which block the phage lysogeny repressor^9^.

In *Bacilli*, many phages have adopted an alternative signaling mechanism, known as arbitrium to control lysis/lysogeny transitions through a peptide-based cell-cell signaling system^10,11^. The arbitrium signaling system is encoded by the phage and consists of the AimR receptor, and the secreted AimP peptide signal. In the absence of the AimP signal, AimR promotes the expression of the *aimX* transcriptional unit through anti-termination^12^. AimP binding to AimR prevents this activity, leading to inactivation of *aimX* expression and establishment of lysogeny^13–15^. In a similar manner to lambda co-infection sensing, the arbitrium system allows phages to become lytic at initial stages of infection when the majority of the bacterial population is uninfected, but switches to lysogeny when a majority of cells are infected^10^. In contrast with lambda co-infection sensing, the arbitrium system has also been shown to function during lysogeny and to regulate lytic induction in concert with DNA damage^16–18^.

The Arbitrium system was mostly studied in the SPβ-like phages ϕ3T and SPβ^10,19^. There, it was found to regulate lysis-lysogeny in association with the phage encoded sroA-F operon and the host encoded MazEF toxin-antitoxin system ^12,17,20–23^. Specifically, in phage ϕ3T, *aimX* was shown to code for a MazE-like antitoxin, AimX^12,20^, while in phage SPβ *aimX* seems to operate at the RNA level, through an unknown mechanism.

Little is known about the mechanisms by which the arbitrium system guides lysis-lysogeny decision in other types of phages and mobile elements. In a study addressing this question, a genomic search identified a diverse phylogeny of AimR genes in *Bacilli*^11^, associated with over a hundred different AimP peptide signals. Arbitrium *aimRP* genes were found to reside on multiple types of phages and on conjugative mobile elements. It was found that in many instances, a small Helix-Turn-Helix (HTH) containing transcription factor homologue was located downstream of the *aimRP* locus in the opposite direction^11^. This led to the hypotheses that: i) the HTH gene codes for the phage repressor and ii) the *aimX* transcript includes the antisense of the repressor transcript and functions to block the transcription or translation of the repressor through sense-antisense RNA interactions. In support of the latter, RNAseq analysis of two such phages during infection, demonstrated that *aimX* transcript extends throughout the putative repressor region and that *aimX* levels are strongly dependent on arbitrium signaling. The same work has also identified the presence of small ORFs in some *aimX* genes and suggested they may function as antirepressors in concert with the antisense regulation. However, neither the nature of the HTH gene, nor the relative importance of the RNA antisense or other regulatory mechanisms have been examined at the molecular level.

Here we show that in a large majority of cases, the arbitrium system is embedded in a conserved five gene locus, including the gene coding for the phage repressor (which we term *aimC*) and a presumably pro-lytic, cro-like, transcription factor, *aimL*. We find that *aimX* is coding for a small antirepressor protein, which blocks the activity of AimC. AimC blocks expression *aimL* and additional putative early lytic genes downstream of it. Finally, by structural, functional and genomic analysis we unveil the molecular mechanism underlying AimX inhibition of AimC.

## Results

### An extended Arbitrium system is found in a majority of phages

To deepen our understanding of arbitrium-coding phages, we constructed a database of arbitrium coding elements (methods). By searching the genomes of ∼5500 group-1 Bacilii (genus: *Bacillus*), we found 9300 AimR homologs in 3800 of these genomes (Fig. 1a, Supp. Fig. 1a, Supplementary File 1). Arbitrium systems often co-resided in the same strain (with an average of 2.4 AimR proteins per strain). Co-residence was even stronger in strains of the *B. cereus* group, where average number of AimR homologs in strains which code for at least one homolog was 2.8, with some strains coding for up to 21 different homologs.

**Figure 1:**
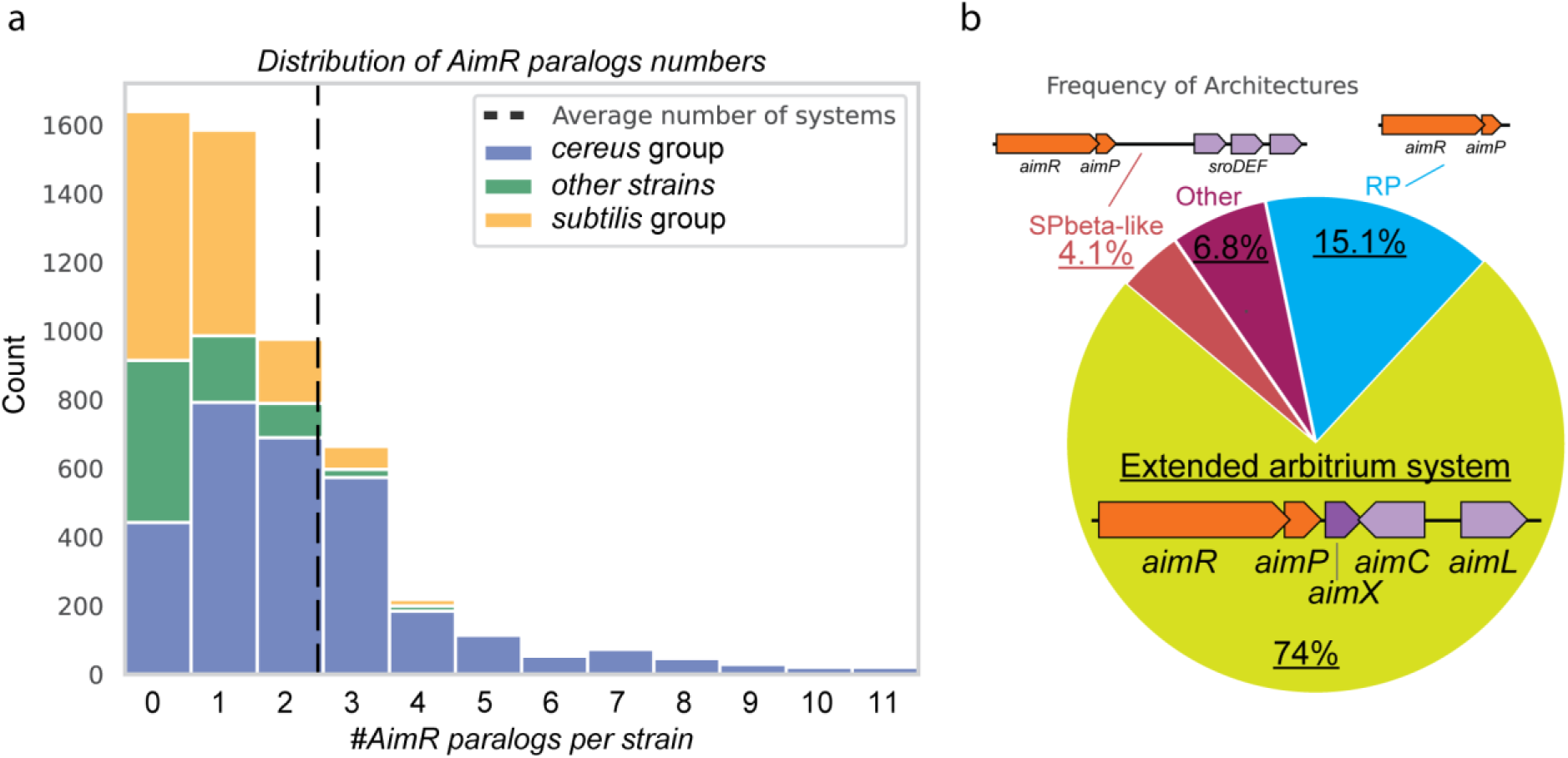
Distribution of arbitrium systems and their genomic organization in *Bacilli*. (a) Distribution of the number of AimR paralogs found in *Bacillus* strains in our database. Different colors mark the distinction between strains from the two main subgroups of this genus (cereus and subtilis groups) and other strains not designated by name to one of these groups. (b) Distribution of architectures in which arbitrium systems are embedded. SPbeta-like and extended arbitrium systems architectures are schematically shown. The “other” slice contains systems where either *aimP* was not found or where either *aimC* or *aimL* genes are found, but not both. The generic RP architecture is marked whenever none of the neighboring genes in either the extended or SPbeta-like systems is found.

We next turned to characterize the genomic context of the *aimR* gene (Fig. 1b, Supp Fig. 1b). Focusing on downstream genes, we first found that 99% of these elements had a clear adjacent putative *aimP* gene. Analyzing further genes downstream of the arbitrium system, we first looked for the well-studied SPbeta-like phages signature by searching for neighboring *sroDEF* genes ^23^. Only ∼4% of the total number of arbitrium systems were identified as part of SPbeta-like phages, all of these systems were associated with AimR of clade 2, as previously identified (Supp. Fig. 2) ^11^. By contrast, in ∼80% of the systems a putative small ORF with a single HTH domain was found immediately downstream and in the opposite direction of the arbitrium genes^11^. We designate this gene *aimC*. 94% of systems coding for such an HTH gene also coded for an additional small HTH domain protein in the opposite direction, which we designate *aimL*. Notably, a homology search for *aimC* and *aimL* yielded very few close homologues not associated with an arbitrium system, further strengthening the direct association of these two genes with the arbitrium system (Supp Fig. 1d).

*aimC* was previously hypothesized to be a repressor gene ^11^, while the *aimC*-*aimL* architecture is reminiscent of the *cI*-*cro* arrangment in phage lambda^5^. We also found that genes downstream of *aimL* and in presumed operonic structure with it were often associated with replication, as in phage lambda (Supp. Fig. 1c). In contrast with phage lambda, the putative AimC proteins were short (80-140aa) and did not code for an auto-cleavage domain. Altogether, our analysis identified that ∼75% of arbitrium systems are embeded in an extended architecture (Fig. 1b).

### aimC represses the aimL promoter, integrating arbitrium and DNA damage control in phage ϕ106

We have previously analyzed the expression of the *aimX* transcript of phage ϕ106, which is found in *Bacillus inaquosorum strain KCTC 13429* ^16^. This phage codes for an extended arbitrium system, with the *aimL* gene followed by several genes in a putative operonic structure, one of which is annotated as coding for a replication organizer protein (Fig. 2a). If the *aimC-aimL* architecture is functionally homologous to the lambda *cI*-*cro* arrangement, we would expect *aimC* to control the *aimL* promoter.

**Figure 2:**
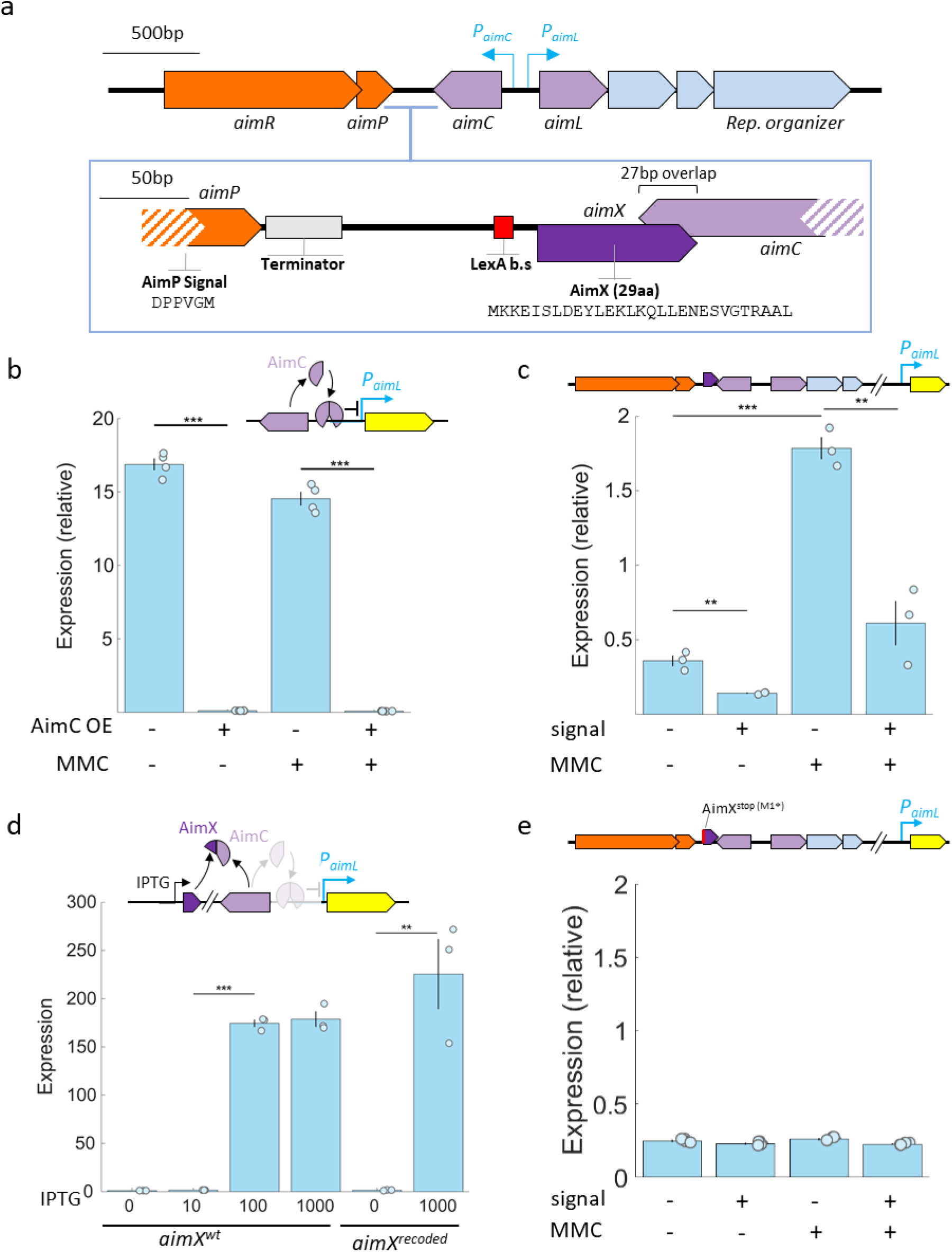
Exploring the regulation of lytic gene expression in the lysogeny module of phage ϕ106. (a) The general architecture of the extended arbitrium locus, showing signaling genes (orange), lysis-lysogeny genes (purple) and early lytic genes (light blue). The region between *aimP* and *aimC* is zoomed-in, emphasizing the *aimP* terminator region, LexA binding site and the putative *aimX* ORF, its protein amino-acid sequence and its overlap with *aimC*. (b-e) Expression from the P*_aimL_*-YFP reporter integrated into an ectopic locus in different strains backgrounds and conditions. (b) Only the reporter or with an additional *aimC* gene and either with or without the addition of MMC. Inset: AimC behaves as a repressor of the *aimL* promoter. (c) Expression in a background encoding for the extended arbitrium locus and two early lytic genes (see inset) in the absence or presence of arbitrium peptide and MMC. (d) Expression in the presence of an *aimC* gene under its own promoter and an IPTG inducible *aimX* gene for varying levels of IPTG (as indicated). Two *aimX* alleles were studied – the wild-type allele (*aimX^wt^*) and a recoded allele (*aimX^recoded^*). Inset: AimX acts as an antirepressor of AimC. (e) as in (c), but for a mutated extended arbitrium system, where the start codon of *aimX* was mutated into a stop codon (*aimX^stop^*). See inset. All experiments were repeated at least three times on separate days. Asterisk mark statistical significance, * p<0.05, ** p<0.005, *** p<0.0005.

To study this transcriptional control, we first cloned a YFP transcriptional reporter of the *aimL* promoter (P*_aimL_*-YFP) and inserted it into a *B. subtilis* PY79 lab strain derivative, deleted for all lytic elements (methods). Flow cytometry was used to measure the median level of fluorescence of the YFP reporter. As we wish to explore the application of DNA damage, we also integrated a constitutive BFP reporter into the strains and used it to normalize for effects related to changes in cell volume during DNA damage. This strain background was then used to probe different parts of the extended system and their response to external conditions (Fig. 2).

First, we compared P*_aimL_*-YFP expression on its own and in the presence of the ϕ106 *aimC* gene expressed under its native promoter. We find that the *aimL* promoter was highly expressed in the absence of the *aimC* gene and that its levels were reduced ∼100 fold in its presence (Fig. 2b), suggesting that *aimC* acts as a repressor. Similar level of repression was found in the presence and absence of the DNA damaging agent Mitomycin C (MMC), suggesting that the short AimC protein indeed does not respond directly to RecA activity (Fig. 2b).

Our previous analysis demonstrated that the *aimX* transcript in this strain is regulated both by AimR and by LexA ^16^. To probe the impact of the arbitrium and the DNA damage control systems on the expression of the *aimL* promoter, we integrated the full extended arbitrium system with two more genes downstream of *aimL* into a second chromosomal locus and measured the expression of the *aimL* reporter with and without the addition of the mature AimP arbitrium peptide (DPPVGM) and the addition of MMC. We find that the *aimL* reporter expression qualitatively matched the previously described expression of the *aimX* reporter ^16^ – expression was activated by DNA damage and repressed by addition of the arbitrium peptide (Fig. 2c).

### Inhibition of aimC by aimX is mediated by a short (29aa) ORF

It was previously suggested that *aimX* regulates *aimC* through antisense RNA (or antisense transcription)^11^. However, further examination of the ϕ106 *aimX* transcript sequence revealed a small ORF of length 29 amino-acids, just downstream of the LexA binding site (Fig. 2a). This raises the possibility that at least part of the regulation of *aimC* is mediated by this short AimX ORF. To test this option, we cloned this ORF under an IPTG-inducible promoter and added this construct to the strain coding for the *aimC* gene and P*_aimL_*-YFP reporter. We assayed the expression of the *aimL* reporter at different levels of IPTG and found that increasing the levels of IPTG led to strong expression of the *aimL* reporter (Fig. 2d). Expression of AimX in the absence of *aimC* had no effect on the *aimL* reporter (Supp. Fig. 2).

In the above construct, *aimX* and *aimC* are expressed from separate loci, preventing *cis* interactions between the two transcripts. However, the AimX transcript has a 30bp overlap in the antisense direction with the end of the *aimC* ORF. This may still lead to *trans* interactions at the mRNA level. To eliminate this interaction, we recoded the overlapping antisense region of AimX to prevent antisense interactions between the two mRNAs (*aimX^recoded^* allele). This allelic change had no significant effect on the ability of AimX to inhibit AimC repression of the *aimL* reporter (Fig. 2d).

While the *aimX* ORF has only a short overlap with the *aimC* ORF, the *aimX* transcript was shown to be further extended in some phages to overlap the full *aimC* ORF antisense. This extended overlap may also have either a cis or trans effect ^11^. To check the importance of the full *aimX* transcript, we replaced the start codon of the *aimX* ORF with a stop codon (*aimX^stop^* allele) within the full arbitrium locus (as assayed in Fig. 2c). We found that the P*_aimL_*-YFP reporter lost its response to either addition of signal or MMC (Fig. 2e). These results support the notion that the *aimX* transcript has little *cis* or *trans* effects on *aimC* expression in the absence of the AimX protein.

### Phage lysis\lysogeny decision is mediated by the AimX antirepressor protein

Our data demonstrate that the arbitrium system controls lysis-lysogeny through protein-level inhibition of the phage repressor. However, we have thus far only probed the level of gene regulation and not its impact on phage decision-making. To this aim, we turned to analyze the role of AimX and the putative repressor within a full phage. We were unable to find a recipient for phage ϕ106 or properly cure it from its host and therefore chose to study the homologous phage ϕW23.

*Bacillus spizizenii* strain W23 was shown to contain a phage designated as ϕW23 ^24^. This phage is related to ϕ106 and codes for a similarly organized extended arbitrium system with a clade 1 arbitrium receptor (Supp Fig 3a,b). While strain W23 is not naturally competent, we were able to induce competence by conjugating into it a plasmid expressing a *comK* gene (methods) ^25^. We used this to construct a phage free strain, termed W23^Δϕ^, and a ϕW23 phage strain marked with an erythromycin resistance gene (designated ϕW23e, methods). Notably, the ϕW23e phage had a lower titer than that of ϕW23 (Supp. Fig. 3c). However, both ϕW23 and ϕW23e were able to infect the W23^Δϕ^ strain and showed similar fold reduction in lysis upon addition of the ϕW23 arbitrium peptide signal (EIIVGA, Fig. 3a, Supp Fig 3c). ϕW23e also showed increased lysogeny upon peptide addition (Supp Fig 3d). We therefore proceeded with the analysis of ϕW23e.

**Figure 3:**
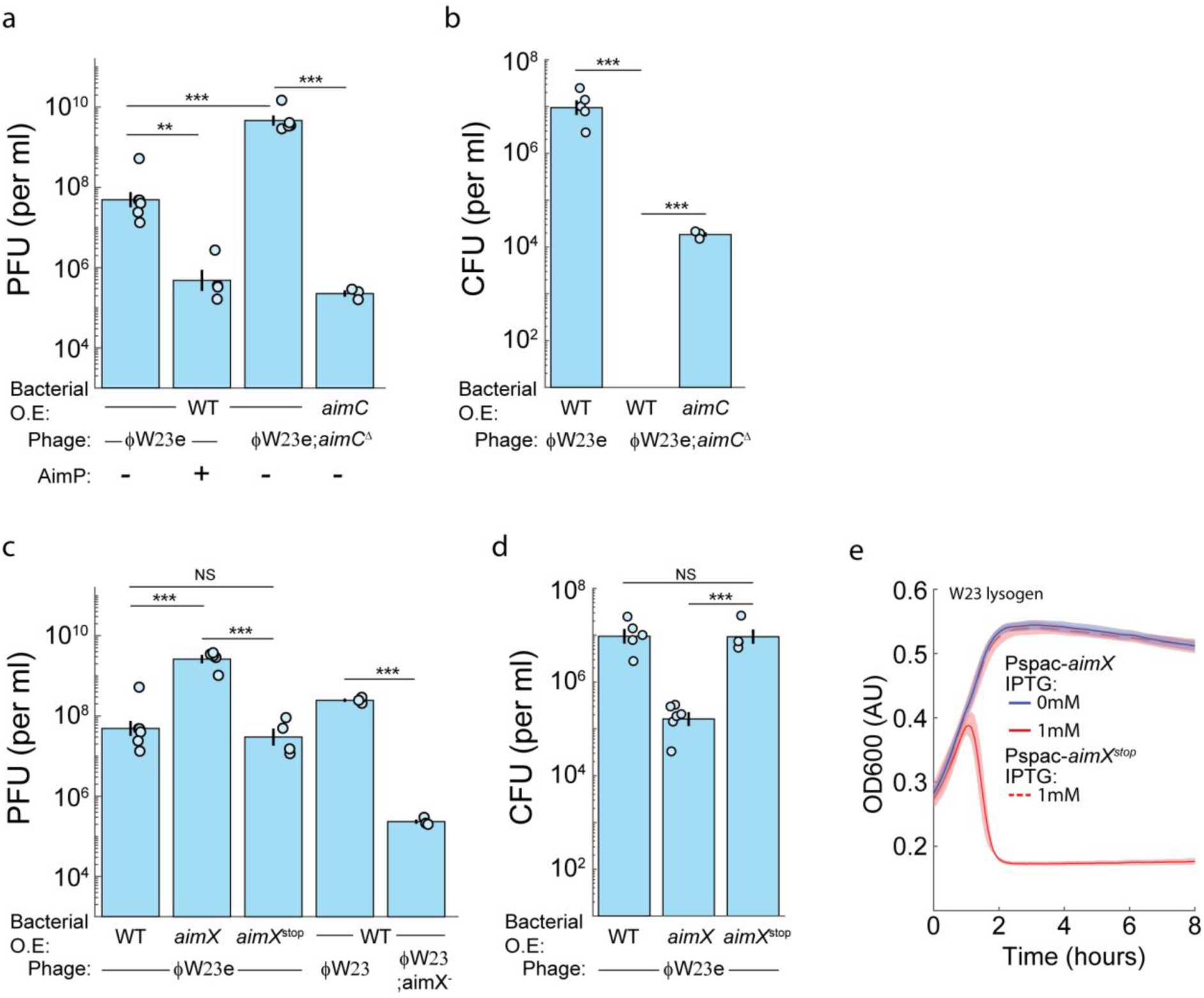
AimC-AimX function as repressor-antirepressor pair in phage ϕW23. Plaque titer (a,c) and lysogen numbers (b,d) measurements done for different ϕW23 and infected cell variants under different conditions, as indicated. Error bars mark standard error, specific data points are marked and where taken on at least three separate days. Asterisks indicate statistical significance, NS – non-significant. (a,b) AimC function as the phage repressor. (c,d) AimX is pro-lytic during infection. (e) AimX is pro-lytic during induction. Induction curves taken with a plate reader, showing cell density upon addition of 1mM (red) or 0mM (blue) of IPTG at time 0, to W23 lysogens coding for an IPTG-inducible *aimX* (solid line) or *aimX^stop^* (dashed line). Both *aimX* transcripts extend (in an antisense direction) throughout the full *aimC* gene. *aimX^stop^*carries a stop mutation at the AimX ORF start codon. Shown are the mean and standard error of at least three measurements taken in different days. All experiments were repeated at least three times on separate days. Asterisk mark statistical significance, * p<0.05, ** p<0.005, *** p<0.0005.

We next probed whether the putative *aimC* repressor gene of this phage indeed acts as the phage repressor. By selecting for free phage particles upon addition of arbitrium peptide, we were able to select for lytic phage mutants in three independent experiments. All sequenced mutants had different mutations in the putative *aimC* gene or its promoter (a C2G substitution at position -55, a single bp deletion at position 213, and a long deletion between positions 265-348). We selected the aimC with the deletion in position 213 (*aimC*^Δ^) mutant for further characterization. We found that the mutant produced no lysogens, but that this was rescued by overexpression of the wild-type *aimC* allele from the bacterial chromosome (Fig. 3a,b). The mutant also showed substantially higher phage titers than the ϕW23e phage. Overexpression of *aimC* led to reduced titer of the *aimC* mutant phage upon infection (Fig. 3a,b). Altogether, these results verify that the putative *aimC* gene is indeed the repressor of ϕW23.

Phage ϕW23 also codes for a putative short ORF within the *aimX* transcript with little homology to the ϕ106 antirepressor (Supp Fig 3B). We wished to determine the relative impact of the AimX_ϕW23_ antirepressor protein and the impact of the *aimC*_ϕW23_ antisense transcript on the lysis/lysogeny decision, during both infection and lytic induction. To this aim, we expressed from the bacterial chromosome under IPTG control either an *aimX ^wt^* allele or an *aimX^stop^*allele. In the latter, the *aimX* start codon was mutated to a stop codon. Both constructs encoded for an extended *aimX* transcript covering the whole antisense of the *aimC* ORF, to allow for a strong sense-antisense mRNA dimerization *in trans*. We found that *aimX*_ϕ_ *^wt^* expression, but not *aimX*_ϕ_ *^stop^* expression was able to induce lysis in the W23 lysogen, to increase the phage titer of an infecting ϕW23::ermR phage and to reduce the number of resulting lysogens (Fig. 3c,d). These results strongly suggest that AimX_ϕW23_ is an antirepressor of AimC_ϕW23_ while the *aimX* mRNA cannot act in trans to regulate phage lysis/lysogeny.

The above results do not rule out a possible *cis* role for antisense transcription of *aimX* on *aimC* expression, e.g, through collision of the RNA polymerase complexes. To explore this, we replaced in the ϕW23 phage the start codon of AimX with a stop codon (methods). We then studied the impact of this ϕW23;*aimX*^stop^ mutation on infection. We found that the mutated phage showed reduced lysis compared to the wild-type phage (Fig. 3d), further suggesting that the *aimX* transcript has little role in lysis/lysogeny decision.

### Putative AimX proteins are found in all extended arbitrium systems

Our results so far point to AimX as a functional antirepressor in ϕ106-family phages, coding for a clade 1 arbitrium system ^11,16^. To characterize the prevalence of AimX antirepressors in other extended arbitrium systems, we searched them for short ORFs downstream of *aimP* and in the same direction. We found such candidates in essentially all these systems. These AimX candidates are typically small (29-80aa), sometimes overlapping the *aimC* gene and are highly divergent, complicating their phylogenetic analysis (Supp. Fig. 4). We performed phylogenetic analysis of the AimC proteins and quantified the level of similarity of the associated AimX proteins within clusters of AimC. The analysis suggest that AimX proteins associated with similar AimC proteins are also similar, though showing larger divergence in their sequence (Fig. 4A, Supp. Fig. 4, Supplementary file 2). This can be expected if AimC and AimX directly interact and co-evolve.

**Figure 4:**
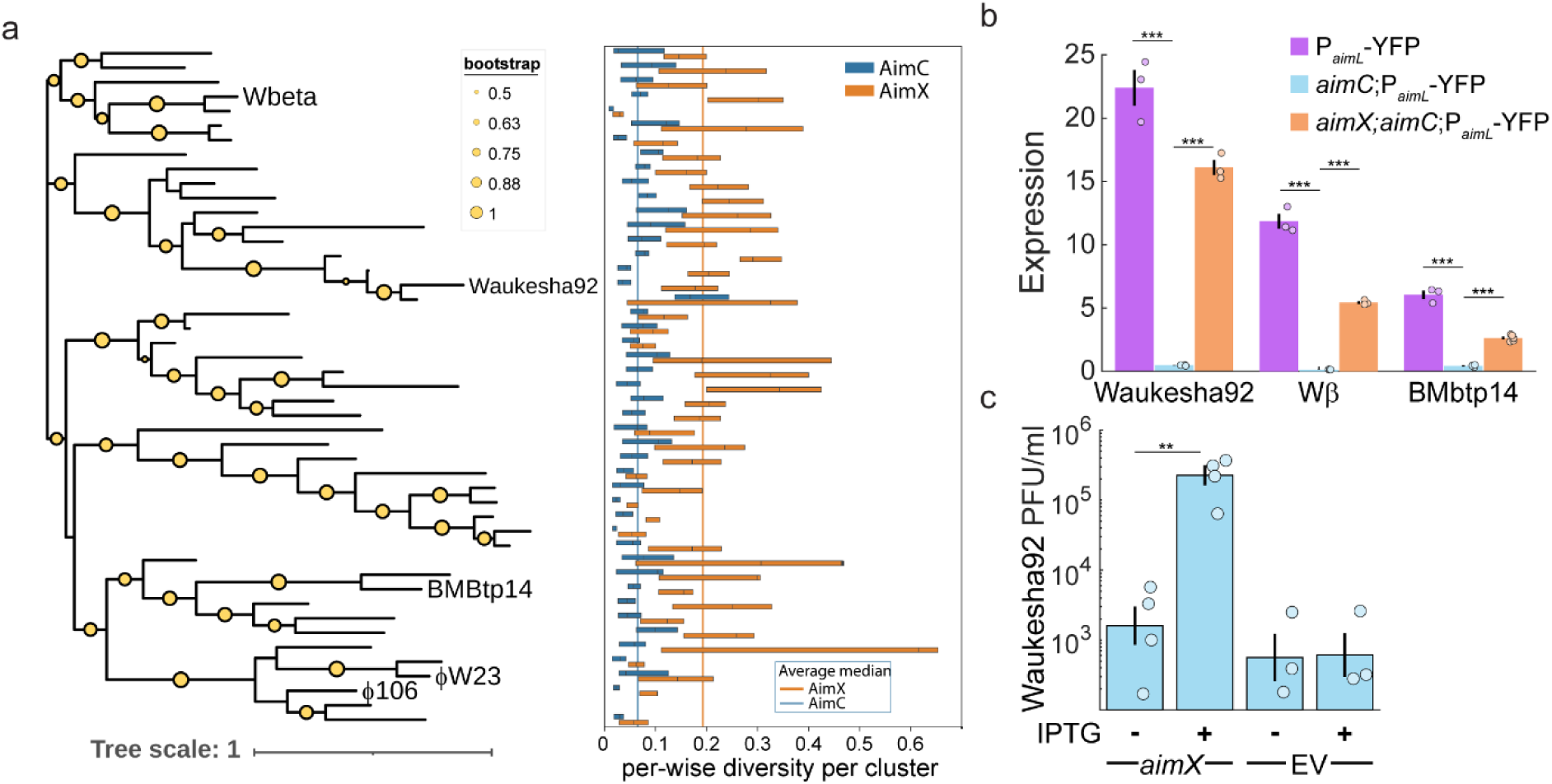
AimX antirepressors are identified in all extended arbitrium systems and verified in additional phages. (a) Left: a phylogenetic tree of AimC proteins. Each leaf is a single representative of a cluster. Clustering of AimC were based on 80% protein identity, as done by DIAMOND^29^. Clusters of specific systems studied in this work are marked. Right: the level of similarity of AimC (blue) and AimX (orange) proteins in each cluster in the phylogenetic tree. Shown are the media similarity and 25^th^-75^th^ percentile values for each cluster and protein based on pairwise similarity between all different variants of the proteins in the same cluster (b) P*_aimL_*-YFP expression (measured in *B. subtilis*) for three different *B. thuriegensis* phages; Waukesha92, Wbeta and BMBtp14. Three different genetic backgrounds were used for each phage – only the *aimL* reporter (purple), reporter and *aimC* repressor expressed from its endogenous promoter (light blue) and *aimL* reporter, *aimC* repressor and IPTG inducible *aimX* antirepressor expressed with 1mM of IPTG (orange). (c) PFU counts after an overnight for strain *B. thuringiensis sv. galleriae BGSC 4G5* carrying the Waukesha92-like phage-plasmid and a plasmid coding for an IPTG inducible AimX or an empty vector (EV), either with or without 1mM of IPTG. All experiments were repeated at least three times on separate days. Asterisk mark statistical significance, * p<0.05, ** p<0.005, *** p<0.0005.

To experimentally verify the generality of the transcriptional repression of the *aimL* promoter by AimC and antirepression by AimX, we studied it in three additional extended arbitrium systems from the *B. cereus* group phages Wbeta, BMbtp14-like and Waukesha92-like (methods) ^11,26–28^. The three phages encode an extended arbitrium system with highly divergent *aimC* repressor and *aimX* genes (Fig. 4A). For each phage, we characterized the expression of its *aimL* promoter by itself, together with the cognate *aimC* gene (with its native promoter) and together with the *aimC* gene and an IPTG inducible cognate AimX. We found that in all cases AimC strongly repressed the *aimL* promoter and that AimX relieved much of this repression (Fig. 4B).

To directly verify the importance of AimX to phage lysis/lysogeny in other phages, we used a phage Waukesha92-like variant (methods) which can infect and lysogenize strain *B. thuringiensis sv. galleriae BGSC 4G5* ^11^. We introduced either an empty plasmid or a plasmid expressing under IPTG control a copy of the phage AimX into this strain and measured titer size after an overnight in both backgrounds with or without IPTG. We find that AimX overexpression led to a 100-fold increase in the number of PFUs (Figs. 4C, methods), supporting the role of AimX as an antirepressor in this phage as well.

### AimX-AimC interaction blocks AimC oligomerization and DNA binding

Our genetic analysis suggests that AimX functions as an antirepressor in a variety of arbitrium-coding phages. However, it is unclear whether this occurs through direct binding and if so, what is the molecular mechanism of antirepression. To answer these questions, we probed directly the interaction of AimX and AimC in two of the phages discussed above, ϕ106 and BMBtp14. We used two methods to probe this interaction. First, we used the *cya* bacterial two-hybrid system to probe binding *in-vivo*^30^. Both AimX_ϕ106_ and AimC_ϕ106_ were fused to the C-terminal of the two *cya* parts (T18 and T25) and their interaction was studied (methods). We found strong homo-dimerization of AimC_ϕ106_ and hetero-dimerization of AimX_ϕ106_ and AimC_ϕ106_, but no evidence for homodimerization of AimX_ϕ106_ (Fig. 5A). Next, we purified these two proteins as well as those of phage BMBtp14. Using SEC-MALS we assayed the molecular mass of the dominant oligomeric state of AimC in solution alone or in the presence of AimX. We find that AimC_ϕ106_ formed a clear dimer in agreement with two-hybrid results, while the mass of the AimC_BMbtp14_ suggested a mixed solution of dimers and tetramers (Fig. 5B). The AimC repressors in presence of the corresponding AimX proteins showed a dominant form of the AimC-AimX heterodimer for both phages (Fig. 5B). Combined, these results suggest that AimX and AimC form a heterodimer that prevents the oligomerization of AimC repressors in both phages.

**Figure 5:**
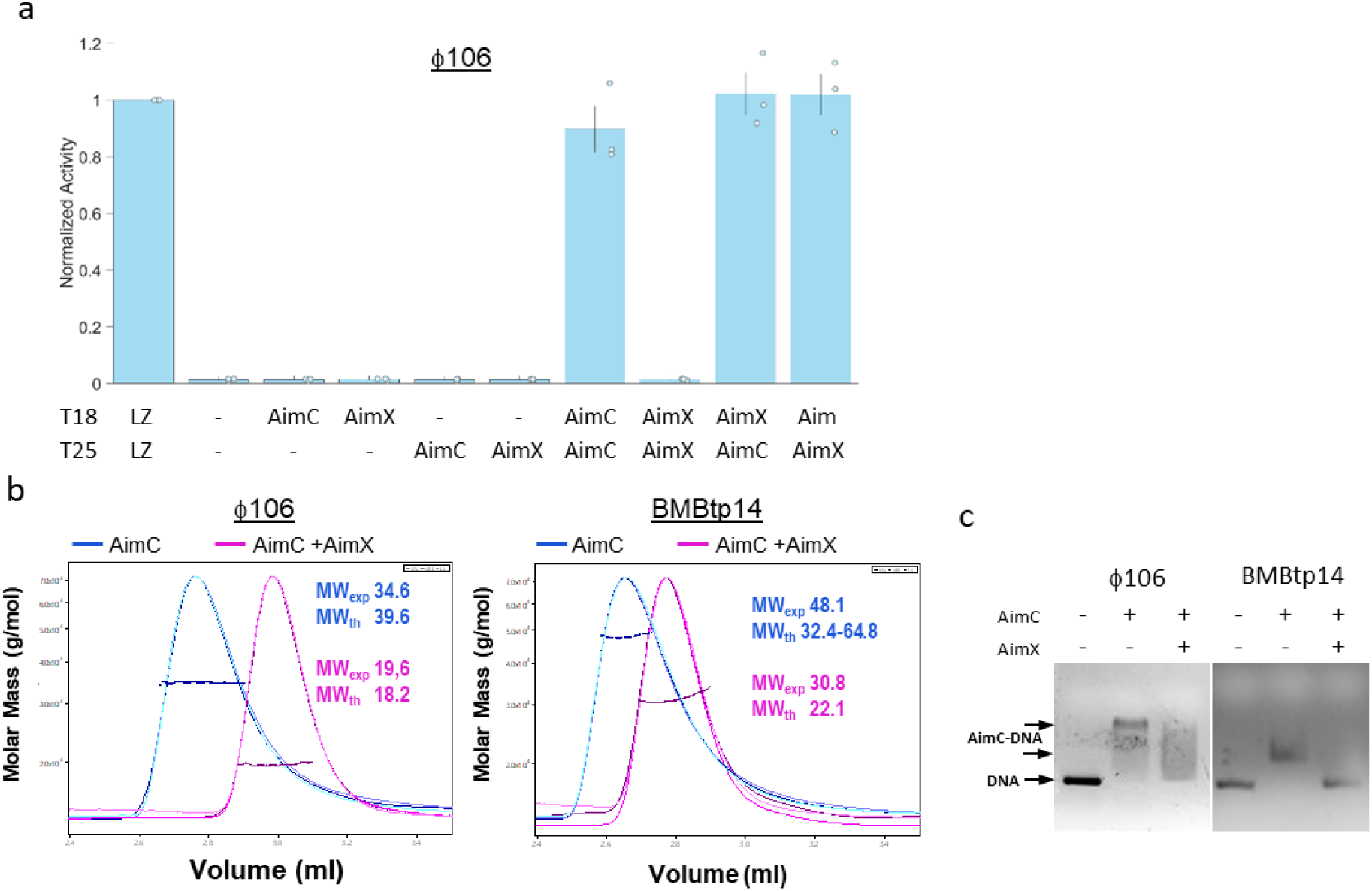
AimX forms a heterodimer with AimC, preventing AimC homodimerization and DNA binding. (a) Bacterial two-hybrid results for interaction between AimX and AimC of ϕ106. Marked are the components fused to the T25 and T18 parts of the Cya protein and the respective mean activity for three independent repeats. Activity in each of three days was normalized by the activity level of the positive Leucine Zipper (LZ) standard control of the assay. (b) SEC-MALS analysis of AimC (magenta) and AimC-AimX (blue) for f106 (left) and BMBtp14 (middle). Chromatograms show the readings from the light scattering (light pink for AimC+AimX and light blue for AimC), refractive index (purple for AimC+AimX and blue for AimC), and UV (dark magenta for AimC+AimX and dark blue for AimC). The horizontal curves represent the calculated molecular masses. Experimental and theoretical molecular weights (MV) for AimC dimers (dimer-tetramer in the case of *aimC*_Btp14_) and AimC-AimX heterocomplex are indicated in kDa. (c) Gel shift experiments (EMSA) after incubation of AimC repressors from ϕ106 and BMBtp14 phages with DNA fragments containing the relatives *aimC*-*aimL* intergenic regions and in presence (1:2 AimC:AimX) or absence of the corresponding AimX antirepressor.

To further show that AimX binding to AimC prevents it from binding to its DNA target, we performed an EMSA DNA binding assay with the whole DNA region between the *aimC* and *aimL* genes of phages ϕ106 and BMBtp14. For both repressors we found that AimC by itself retarded DNA migration, suggesting that it binds to this DNA fragment. In contrast, when AimX was added together with AimC, the retarded band became weaker or disappeared, indicating that the interaction with AimX prevented AimC from binding to its operator (Fig. 5C).

To explore the mechanism of antirepression we combined structural data and modeling. We were able to obtain the crystal structure of AimC_BMbtp14_ at 1.9 Å resolution (Supp.Table Val1) showing that AimC_BMbtp14_ is composed of two folded domains connected by an unstructured linker of ten residues (94-103) (Fig. 6A). The N-terminal domain (residues 19-93) presents the prototypical helix-turn-helix (HTH) fold of DNA binding proteins composed of five (α1-α5) helices. The C-terminal domain (residues 104-142) is composed of two antiparallel helices (α6 and α7) connected by a short β-hairpin (β1-β2) disposed perpendicular to the helices (Fig. 6A and Supp Fig. 5). This small domain seems to present a new fold since search with Foldseek ^31^ and Dali ^32^ servers did not yield results with significant scores. In the crystal structure, the asymmetric unit contains four AimC_BMbtp14_ monomers organized in two dimers that form a dimer of dimers (Supp. Table 1 and Supp. Fig. 5). Repressor oligomerization is mediated almost exclusively by the C-terminal domains, with a larger interaction surface (∼1100 Å^2^) mediating dimer formation and a smaller tetramerization surface (∼390 Å^2^) provided exclusively by α7 helix (Fig. 6A, 6C and Supp Fig. 5). This structural oligomeric organization fits well with the combined dimer/tetramer species found in the SEC-MALS data (Fig. 5B).

**Figure 6:**
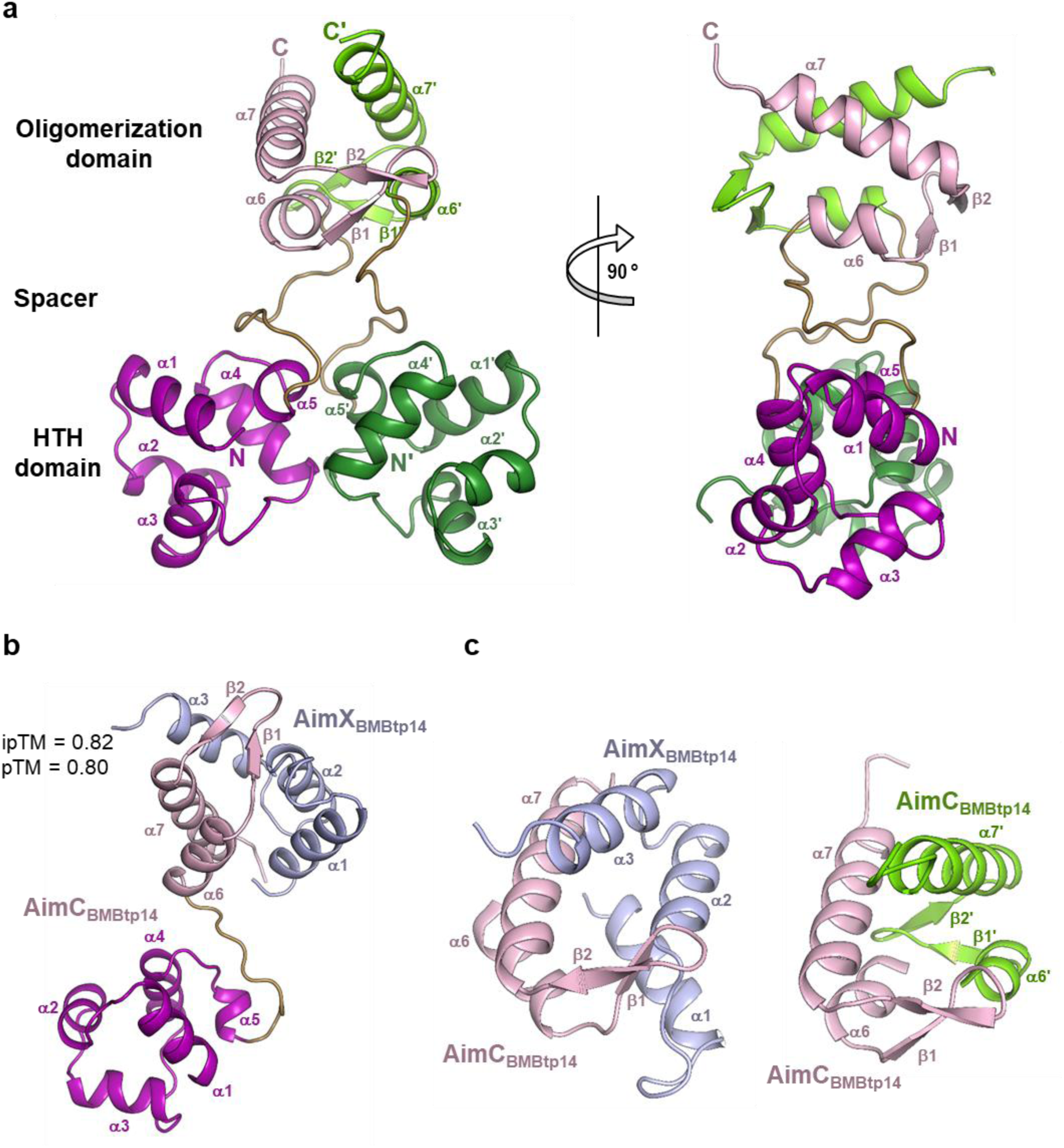
**Structural characterization of** *aimC*_BMbtp14_ and its complex with AimX_BMbtp14_. a) Two orthogonal views of the ribbon representation from the AimC_BMbtp14_ homodimer in the crystal structure. Each subunit is coloured in purple and green with the HTH and oligomerization domains in dark and light tones, respectively. Structural elements are labelled (‘ denotes element in the second subunit). b) Alphafold 3 model of AimC_BMbtp14_-AimX_BMbtp14_ heterodimer in ribbon representation. AimC_BMbtp14_ is coloured as in A and AimX_BMbtp14_ is coloured in light-blue. Structural elements are labelled. c) Close view of the C-terminal oligomerization domain of AimC_BMbtp14_ in the heterodimer with AimX_BMbtp14_ (left) or in the AimC_BMbtp14_ homodimer.

We were unable to crystalize AimC_BMbtp14_ in complex with AimX_BMbtp14_. Therefore, we used AlphaFold 3 ^33^ to predict the putative structure of the AimC_BMbtp14_-AimX_BMbtp14_ heterodimer observed in the SEC-MALS assays. AimX_BMbtp14_ is predicted to adopt a three-helix fold and bind the AimC_BMbtp14_ C-terminal oligomerization domain, covering the dimerization and tetramerization surfaces of the AimC_BMbtp14_ monomer (Fig 6B and 6C). We also analysed the AimX-AimC Alphafold binding models for additional AimC repressors tested in this work (ϕ106, Wbeta and Waukesha92). The model predicts with good scores a dimer organization for ϕ106 and Wbeta and a tetramer organization for Waukesha92. These were similar to the AimC_BMbtp14_ structure, with an N-terminal HTH domain followed by a variable unstructured connector and a C-terminal domain with alpha helices, mediating much of the oligomerization (Supp. Fig. 6). Modelling of the heterocomplexes with the corresponding AimX shows that in all cases the anti-repressor adopts a predominant alpha-helix fold (a β sheet is predicted only for AimX_ϕ106_), but reduced to one or two helices. Alphafold predicts with high confidence (ipTM score higher than 0.8) that the AimXs interact with the C-terminal oligomerization domains of the corresponding repressor and occupies the position of the second C-terminal domain of the repressor in the AimC dimer (Supp. Fig. 6). Modeling therefore suggests that the mechanism of action of AimX is conserved, interacting with the AimC C-terminal domain and preventing repressor oligomerization required for DNA recognition and binding. Since the repressors show variability in their C-terminal domains, AimX structure also varies although helical folding prevails. This difference on AimC-AimX structures, suggest that AimX binding to AimC may be specific.

## Discussion

In this work we lay out the mechanism underlying lysis/lysogeny decision in the large majority of arbitrium-coding systems. We identified an extended core arbitrium module containing 5 genes. In addition to the arbitrium receptor, AimR, and signaling gene, AimP, which have been previously identified, the system codes for a small antirepressor, AimX, AimC which serves as the phage repressor and for AimL; most likely a *cro*-like gene whose function remains to be explored. In the absence of AimP, AimR enables the expression of AimX, which blocks the AimC repressor. This activates the lytic program by allowing the expression of the early lytic AimL operon. When AimP concentration is high, AimR is inactive and so the lytic program is not activated. Our analysis suggests that the previously posited sense-antisense interaction at the RNA level between the AimX transcript and AimC is most likely not consequential to arbitrium’s function^11^.

We also found most arbitrium systems we have analyzed to be activated by DNA damage. In ϕ106-family phages this is regulated through LexA binding to a specific site upstream of *aimX* (Fig. 2a, Supp. Fig. 3b). However, many of the phages with an extended arbitrium system do not show such a LexA binding site upstream of *aimX.* Further work is required to elucidate the mode of regulation by DNA damage in various extended arbitrium systems.

We note that the regulator of lysis is called AimX in all arbitrium phages, but that in fact there are currently three known different mechanism by which *aimX* functions. In SPβ, *aimX* seems to function as an RNA as originally proposed^10^, though the way by which it works is still unknown. In phage ϕ3T, AimX codes for a MazE-like antitoxin that blocks the activity of the bacterial MazF toxin ^12,20^. Finally, in the extended systems discussed here, AimX functions as the antirepressor of the phage master repressor, AimC. It is interesting that the same design criterion is implemented so differently in different phages exploiting identical communication mechanism. The advantages and disadvantages of these different designs is yet unclear. Future works should take care in distinguishing these genes which share a common name and general function, but vary greatly in the mechanism of action.

At the molecular level, the interaction between the AimX antirepressor and the the AimC repressor is direct. AimX binding to AimC prevents its oligomerization and consequently its binding to the repressor operator sites. This AimX molecular mechanism of antirrepression is reminiscent of the well-studied interactions between the SinR *Bacillus* repressor of biofilm formation and its SinI antirepressor, where the small helical SinI antirepressor mimics the oligomerization domain of SinR^34,35^. SinI interaction produces a SinI-SinR heterocomplex and precludes the SinR tetramerization required for DNA biding in a similar way as the AimC_BMbtp14_ structure and AimC_BMbtp14_-AimX_BMbtp14_ model proposes.

Small phage protein are highly abundant and their role in anti-defense, host modulation and other phage functions is actively explored ^36,37^. Recent works have shown the extensive use of small antirepressors (of order 80aa) in the control of phage lysis-lysogeny in *Vibrio* phages ^38^. Our work here reinforces and expands on this growing group, by unveiling a novel system of antirepression that depend on very small proteins (down to 29aa long). Such small proteins are often hard to identify, especially if they overlap other open reading frames, as occur in our case. Our work therefore contributes to the ongoing challenge to identify the extent of small protein regulation and function in phage biology ^36^.

**Table 1.**
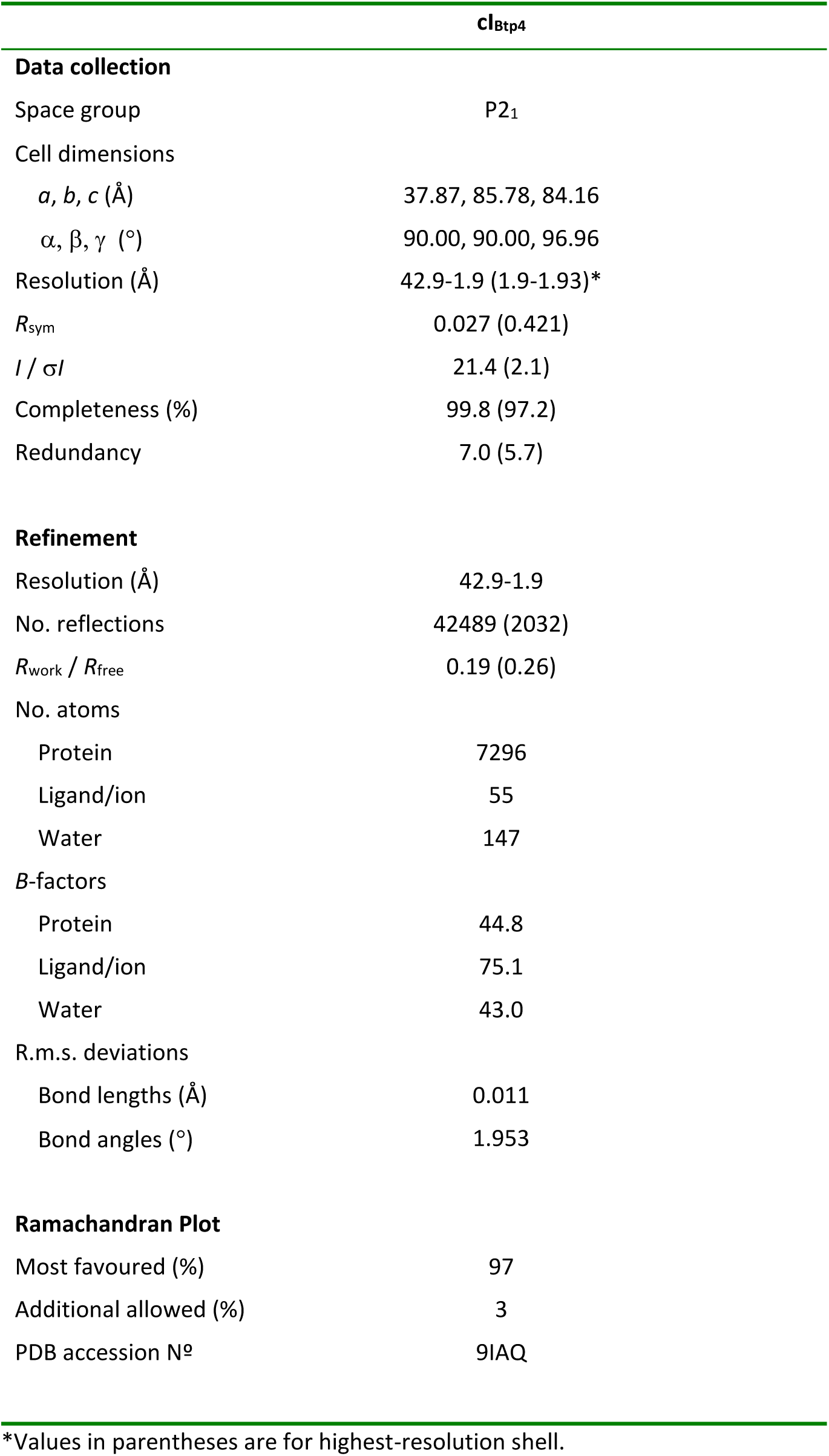
Data collection and refinement statistics for protein structure.

## Acknowledgment

We thank Zhihui Xu for providing us the pUBXC plasmid ^25^ and Annika Gillis and her group for help with protocols and plasmids for working on *B. cereus* group strains. We thank Nitzan Aframian and other members of the Eldar lab for comments on the manuscript as well as Jose Penades and Cora Chmielowska. We thank Marta Segade Vázquez for her help with AimC_Bmbtp14_ and AimX_Bmbtp14_ production and purification. We thank the IBV-CSIC Crystallogenesis Facility for protein crystallization screenings. Data collection experiments for structural data were carried out at XALOC beamline at ALBA synchrotron (Cerdanyola del Valles, Spain). X-ray diffraction data collection was supported by block allocation group (BAG) ALBA Proposal 2023077633. We acknowledge the ALBA synchrotron for provision of beam time and we would like to thank beamline staff for assistance. This work was supported by grants No. 2288/2021 from the Israel Science Foundation to A.E.; PID2022-137201NB-I00 from Spanish Government (Ministerio de Ciencia e Innovación), CIPROM/2023/30 from Valencian Government and the European Commission NextGenerationEU fund (EU 2020/2094), through CSIC’s Global Health Platform (PTI Salud Global) to A.M. A.E. and A.M. are funded by European Research Council Grant 101118890 (TalkingPhages).

## Materials and Methods

### Constructing the Arbitrium Database

We conducted a comprehensive survey of 5,491 genomes belonging to Bacillus rRNA group 1, all obtained from the National Center for Biotechnology Information (NCBI) database. To identify Arbitrium systems across these genomes, we implemented a multi-step approach targeting the core components of the system. We searched for AimR homologos by running tblastn locally^39^ with an E-value threshold of 1e-5. To perform this search, we assembled a database of AimR sequences that had already been classified into their respective AimR clades ^11^. For each identified AimR homolog, we examined the genomic region 250 bp downstream of the AimR coding sequence (allowing for minor overlaps) to identify putative AimP sequences. We identified candidates by screening predicted short ORFs in this region. These putative AimP sequences were then validated by comparing them to an established list of known AimP signals and also by confirming their function as signal peptides using SignalP 5.0 ^40^. We searched for putative AimX sequences in the region downstream of confirmed AimP sequences. Candidate AimX sequences were filtered based on length (<75 a.a) and their predicted secondary structure characteristics, specifically looking for a high degree of alpha helicity. Secondary structure predictions were generated using the Jpred 4 webserver^41^. Putative AimC and AimL sequences were identified based on their genomic positions relative to the Arbitrium system components. The AimC candidates were located downstream of AimX on the strand opposite to AimR, while AimL candidates were located in the same genomic region but immediately downstream of AimC and on the same strand as AimR. Both proteins were verified by the presence of helix-turn-helix (HTH) DNA-binding motifs.

Annotation beyond the Arbitrium locus was achieved by predicting and annotating all ORFs found within 50 kbp upstream and downstream of each identified AimR homolog. First, we ran Pharokka^42^ to identify potential phage genes and their functions. We then applied Phold (https://github.com/gbouras13/phold) for improved functional annotation, which uses protein structural homology to enhance annotation sensitivity^43,44^.

### whole genome comparison of ϕW23 and ϕ106

A visualization of the whole genome comparison of the two phages was created with Clinker ^45^ using the default parameters (identity threshold of 0.3).

### AimC and AimL pair search

HMM profiles of AimC and AimL were constructed with HMMER v3.4 (http://hmmer.org) using the ’hmmbuild’ command with default parameters from manually curated multiple sequence alignments. These profiles were then searched against our database of Bacillus rRNA group 1 genomes, using the ’hmmsearch’ command with an E-value threshold of 1e-5. A custom Python script was used to parse the results and identify any AimC and AimL pairs located adjacent to each other (within 500bp) and in opposite orientations (one on the forward strand, one on the reverse). The script also determined whether these pairs were found near an AimR gene (≤ 10 ORFs away in either direction). For cases where a pair was identified within 10 ORFs of a contig boundary, the pair was excluded from downstream analysis to avoid potential false negatives in AimR detection due to genome fragmentation. All of the identified AimC and AimL were then clusters using DIAMOND ^29^. The ‘cluster’ command was executed with the --approx-id parameter set to 80%.

### Phylogenetic analysis and clustering of AimC and AimX

The protein sequences of AimX and AimC were clustered using DIAMOND ^29^. The ’cluster’ command was executed with the --approx-id parameter set to 80%. A total of 47 AimC clusters were selected for downstream analysis based on their size, with the exception of clusters 46 and 47, which were included because the AimC sequences of ɸ106 and BMBtp14 were assigned to them. For each of the clusters a representative sequence was chosen. All of those sequences were multiply aligned using MAFFT ^46^, and a phylogenetic tree was generated using FastTree^47^. The tree was visualized using iTOL^48^. To assess sequence diversity within each cluster the pairwise sequence identity was computed using global alignment with Biopython’s PairwiseAligner^49^. Sequence identity was defined as the fraction of identical residues in the shortest sequence length, and sequence distances were calculated as 1−identity. For each cluster, we calculated the distances between all the unique sequences of AimX and AimC. Statistical summaries of these distances were compiled separately for AimX and AimC sequences and visualized using a boxplot.

### Growth media and conditions

All the experiments in this study were performed in rich media- Luria Bertani Broth (LB): 1% tryptone (Difco), 0.5% yeast extract (Difco), 0.5% NaCl. Liquid cultures were grown with shaking at 220 RPM and a temperature of 37°. When preparing plates, medium was solidified by addition of 2% agar. Antibiotics were added (when necessary) at the following concentrations: ampicillin: 100 µg ml^-1^, spectinomycin: 100 µg ml^-1^, chloramphenicol: 5 µg ml^-1^, kanamycin: 10 µg ml^-1^, erythromycin 3 µg ml^-1^, zeocin10 µg ml^-1^. For experiments that involve infection, LB was supplemented with 0.1 mM MnCl_2_ and 5 mM MgCl_2_ to create MMB media. The P_hyperspank_ and P_spac_ promoters were activated using 1 mM isopropyl-ß-D-thiogalactopyranoside (IPTG) unless stated otherwise, and the P_man_ promoter was activated using 0.2% mannose. *B. thuringiensis* strains containing pBHE plasmid were grown at 30° with chloramphenicol to ensure plasmid maintenance.

### Strain & plasmid construction

All bacterial strains, plasmids, and primers used in this study are listed in Supplementary file 3. To construct *B. subtilis* strains, standard transformation protocols were used for genomic integration and plasmid transformation ^50^. The *B. spizizenii strain W23* was transformed by inducing a xylose inducible comK from the pUBXC plasmid, as previously described ^25^. *B. thuringiensis* strains were transformed via conjugation with E.coli SM10(λpir) containing the plasmids of interest.

All plasmids were constructed by restriction and ligation or using Gibson assembly. Marked ϕW23 was constructed by insertion of an erythromycin resistance cassette between the 09480 and 09495 genes using a long flanking homology PCR method. For deletion of the muramidase gene in ϕW23 prior to curing, we used a long flanking homology PCR method to replace the gene with a kanamycin resistance cassette.

Marked Waukesha92-like phage-plasmid was constructed by insertion of a spectinomycin resistance cassette between the 34651 and 34656 genes. Briefly, a spectinomycin cassette and upstream and downstream regions of the cassette insertion locus were amplified and cloned into the pBHE vector (pKSV-oriT) by Gibson assembly. The plasmid was then conjugated to *B. thuringiensis sv. galleriae BGSC 4G5 Waukesha-92 like lysogen* using the *E. coli* SM-10 strain. *Trans-*conjugants were selected on LB agar plates supplemented with chloramphenicol (for the plasmid) and Polymyxin B Selective supplement (for gram-positive bacteria) (Merck **P9602**) and transferred to LB plates supplemented with chloramphenicol at 42 °C, to promote plasmid integration into the bacterial chromosome by homologous recombination. The bacteria were then passed several times in fresh LB medium without chloramphenicol at 37 °C to promote curing of the plasmid. Single colonies were screened by plating on LB plates with or without chloramphenicol to confirm plasmid curing. Chloramphenicol-sensitive colonies were then tested for spectinomycin cassette integration by PCR.

Plasmids for protein expression were generated using *NEBuilder HiFi Assembly* protocol. Genes for cloning were obtained by synthesis with sequence optimized for expression in *E. coli* (Table Val1) including overlapping ends for the assembly in the plasmids. Plasmid pETduet was used as template for plasmid amplification using the corresponding primers indicated in supplementary file 3. *aimC*_BMbtp14_ and *aimC*_ϕ106_ were cloned at pETduet site 1 with an N-terminal His-tag followed by a TEV protease cleaving site. For *aimC*_BMbtp14,_ the full length ORF annotated in phage genome (YP_009830673.1) was cloned, although the 17 first aminoacids are not present in other members of AimC family and an alternatively V18 could be used as start codon. In the case of AimC-AimX coexpression, *aimX*_BMbtp14_ and *aimX*_ϕ106_ were cloned at site 2 of pETduet plasmids with the corresponding AimCs including an N-terminal S-tag.

### Plaque forming assay

Samples for PFU measurements were collected from cultures centrifuged for 5 min at 4,000 rpm at room temperature. Next, the supernatant was filtered using a 0.22 μm filter (Sartorius biotech cat. 14-555-270). 100 μL of filtered supernatant at an appropriate dilution were then mixed with 200 μL of the relevant *B. subtilis* strain grown in MMB to to OD_600_ of 0.3, and left to incubate at room temperature for 5 minutes. 3 ml of molten MMB with 0.5% agar medium (at 60°C) were added, mixed, and then quickly overlaid on LB-agar plates. Plates were then incubated at 37° for 1 hour following an overnight incubation at room temperature, to allow plaques to form.

### Induction experiments

Strains were grown overnight in LB at 37° with shaking at 220 RPM then diluted by a factor of 1:100 into fresh LB medium. Upon reaching OD_600_ = 0.3 cultures were supplemented with 1mM IPTG, when indicated.

Optical density measurements at a wavelength of 600 nm were performed in a 96-well plate using a plate reader (INFINITE® multimode microplate reader, Tecan, USA).

### ϕW23 Infection experiments

Strains were grown overnight in LB at 37° with shaking at 220 RPM then diluted by a factor of 1:100 into fresh LB media. Upon reaching OD_600_ = 0.3 cultures were supplemented with 1mM IPTG, 10 µM of EIIVGA peptide, when indicates. ϕW23 phage was added at a MOI of 0.1. For quantification of lysogens, 1ml of infected cultures were taken 30 minutes or 2 hours after infection. Cells were washed with 1 ml of PBS to remove unbound phage particles and then diluted and spread on LB plates with eryrhromycin to select for cells that had become lysogenized.

For PFU quantification, 1 ml of infected cultures was collected 2 hours after infection, centrifuged for 5 min at 4,000 r.p.m at room temperature, and filtered using a 0.22 μm filter. Then, a plaque forming assay was performed.

### Waukesha92-like experiments

Strains were grown overnight in LB supplemented with chloramphenicol 5 µg ml^-1^ and spectinomycin 200 µg ml^-1^ at 30° with shaking at 220 RPM. They were then diluted by a factor of 1:100 into fresh LB media containing chloramphenicol and spectinomycin, with or without 1mM IPTG, and incubated at 30° with shaking at 220 RPM. ON cultures were centrifuged for 5 min at 4,000 rpm at room temperature and the supernatant was filtered using a 0.22 μm filter. 100 μL of filtered supernatant at an appropriate dilution were then mixed with 200 μL of *B. thuringiensis sv. galleriae BGSC 4G5 cured* strain grown in MMB to to OD_600_ of 0.3, and left to incubate at room temperature for 5 minutes. 3 ml of molten MMB with 0.5% agar medium (at 60°C) were added, mixed, and then quickly overlaid on LB-agar plates. Plates were then incubated overnight incubation at 30°, to allow plaques to form.

### Phage ϕW23 Curing

Phage curing was performed using a ϕW23 lysogenic strain harboring an IPTG-inducible *aimX* gene and a deletion of the muramidase gene. The strain was plated on LB agar supplemented with IPTG to induce *aimX*, triggering prophage induction and subsequent cell lysis. After incubation, only *Bacillus* cells that had lost the prophage were able to form colonies. To ensure that the phage cured cells are not reinfected by phages released from lysed cells on the plates, the muramidase gene was deleted rendering the phage unable to infect.

#### Flow cytometer and tracking gene expression by fluorescent reporters

Flow cytometry was performed to quantify gene expression at the single-cell level, using a Beckman-Coulter Cytoflex flow-cytometer equipped with four lasers (405 nm, 488 nm co-linear with 561 nm, 638 nm). The emission filters used were: BFP – 450/50, YFP – 525/40. Constitutive mTag2-BFP was used to distinguish the DNA damage effect on the cells and the treatment effect by dividing the mean YFP expression with the mean BFP expression.

To determine expression, cultures were grown in LB overnight at 37° with shaking at 220 RPM and then diluted by a factor of 1:1000 into fresh LB media containing different concentrations of IPTG or/and 2% xylose, as indicated. Upon reaching O.D600 = 0.3, 10 µM of the appropriate arbitrium peptide (DPPVGM for ɸ106; EIIVGA for ɸW23) or/and 0.5 µg ml-1 of MMC were added. The YFP and BFP expression levels were measured using flow cytometry at different time points.

#### Bacterial two-hybrid

Cells were grown overnight at 30° with shaking at 220 RPM in Acumedia^©^LB supplemented with 100 ug mL-1 ampicillin, 30 ug mL-1 kanamycin and 0.5mM IPTG. Β-galactosidase activity was measured using the ONPG assay in a 96-well array in a TECAN plate reader, as previously described, and calculated as ((OD420nm at time t2–OD420nm at time t1)/t2–t1 (min))/OD600nm. The t2 and t1 time points are chosen to be located in the linear part of the kinetic.

### Protein expression and purification

All proteins were produced and purified according to described methods ^13,51^. Briefly, single colonies carrying the corresponding expression plasmid was grown overnight at 37°C in 100 ml of LB medium supplemented with 100ug/ml of ampicillin and 33ug/ml of chloramphenicol. A 1:100 dilution of the overnight culture was used to inoculate 4L of autoinducing medium containing ampicillin and chloramphenicol. The culture was grown at 37°C with agitation. When OD_600_ reached 0.9 the temperature was set to 20°C and incubated for additional 18h. Cells were harvested by centrifugation at 3000 *g* for 40 minutes and the pellet was suspended in lysis buffer (50 mM TRIS pH 8, 500 mM NaCl, 1 mM MgCl_2_ for AimC_ϕ106_, AimC_ϕ106_-AimX_ϕ106_ and AimC_Bmbtp14_-AimX _Bmbtp14_ or 50 mM Hepes pH 7, 500 mM NaCl, 1 mM MgCl_2_ for AimC_Bmbtp14_) and lysed by sonication on ice. Lysate was centrifuged at 10000 *g* for 1h to remove cell debris and the supernatant was loaded on a 5ml HisTrap FF (Cytiva), washed with 5 column volumes of lysis buffer supplemented with 10mM imidazole and eluted with lysis buffer supplemented with 500 mM imidazole. The fractions containing the proteins of interest were collected and digested with TEV protease (protein:TEV molar ratio 50:1) while they were dialyzed against lysis buffer supplemented with 0.5 mM EDTA and 1 mM β-mercaptoethanol. Dialyzed samples were loaded on a 5ml HisTrap FF (Cytiva) to remove the His-tag and the protease, and the fraction not retained in the column containing the His-tag free protein was collected and concentrated by centrifugation through an Amicon Ultra system (3 kDa cutoff) and loaded in a Hi-Load Superdex 75 10/300 (GE Healthcare) equilibrated in lysis buffer. The eluted fractions were judged by SDS-PAGE and the those with the purest protein were pooled, concentrated and stored at -80°C.

### Protein Crystallization and data collection

Crystals from AimC_BMbtp14_ were grown in sitting drops at 294°K using vapor-diffusion method. Initial crystallization trials were set up in the Cristalogenesis service of the IBV-CSIC using commercial screens JBS I, JBS II (JENA Biosciences) and JCSG (Molecular Dimensions) in 96-well plates (Swissci MRC2) using equal volumes of protein at 10 mg/mL and mother liquor. Crystals grew in 1 day using 10% PEG 8000, 10% ethylene glycol and 100 mM HEPES pH 7.5 as mother liquor. Crystals were flash frozen in liquid nitrogen using mother liquor supplemented with 20% glycerol as cryoprotectant. Diffraction data was collected at 100° K from single crystals at XALOC beamline of ALBA Synchrotron. Data sets were processed using XDS ^52^ and merged and reduced with Aimless ^53^ in CCP4 suite ^54^. The data-collection statistics for data sets used in structure determination are shown in Supplementary Table 1.

### Phase determination, model building and refinement

AimC_BMbtp14_ crystal structure was solved by molecular replacement with MOLREP ^52^ using the coordinates of the DNA binding domain of SinR (PDB 3QQ6) as search model. The electron density maps calculated with these phases were of sufficient quality to manual building of the C-terminal oligomerization domain of the protein with COOT ^55^. The final model was refined by alternating several rounds of manual building and automatic refinement with REFMAC ^56^. The final AimC_BMbtp14_ model includes all residues of the expressed ORF with the exception of the initial 17 residues corresponding to a possible additional region due to a discrepancy by the start codon. This region is not visible in the density map indicating that it is highly flexible and that most likely the natural start codon corresponds to Val18 in the ORF. Refinement statistics are summarized in Supplementary Table 1. Figures for structural representations were generated with Pymol software (https://pymol.org/2;Schrödinger).

### Protein model prediction

The candidate sequences of AimC repressors and AimX antirepressors from phages BMBtp14, ϕ106, Wbeta and Waukesha92 were submitted to AlphaFold 3 server^33^ (https://alphafoldserver.com) and the model for AimC dimers or tetrames and AimC-AimX heterodimers were calculated using standard setups. Predicted structures were visualized and figures were generated using PyMOL (Schrodinger Inc.).

### EMSA assays

The ability of AimC_ϕ106_ and AimC_BMbtp14_ to bind the DNA located between *aimC* and *aimL* genes of both phages was analyzed by native agarose gel electrophoresis. Probes were obtained by PCR using primers summarized in Supplementary File 3. Fragments were purified using GeneJet PCR Purification Kit (ThermoFisher®). DNA binding reactions were performed for 5 min at room temperature in a volume of 20 μl containing 15 μg de DNA and 7.5 μM of AimX repressor. The effect of AimX_ϕ106_ or AimX_BMbtp14_ on DNA binding was evaluated by adding (15 μM final concentration) at the corresponding AimC-DNA mixture and the incubation for 5 additional min at room temperature. Electrophoresis was then performed in 1.5 % agarose gels in Tris-Acetate-EDTA (TAE) buffer for about 40 min at 80V at room temperature.

### Size Exclusion Chromatography with Multi-Angle Light Scattering (SEC-MALS)

A Wyatt DAWN HELEOS-II MALS instrument and a Wyatt Optilab rEX differential refractometer (Wyatt) coupled to a Shimadzu HPLC were used to perform SEC-MALS experiments. 20 μl of AimC or AimC-AimX (1:1 molar ratio) at 2.5 mg ml^-1^ were injected in a KW-803 (Shodex) column previously equilibrated in 25 mM Hepes pH 7, 250 mM NaCl and were eluted isocratically in the same buffer at a flow rate of 0.3 ml min^-1^. The Astra 7.1.2 software provided by the manufacturer was used for the acquisition and analysis of the data.

### Structural Data Availability

Coordinates for atomic structures have been deposited at the RCSB Protein Data Bank (PDB code 9IAQ).

---

**Supplementary Figure 1:**
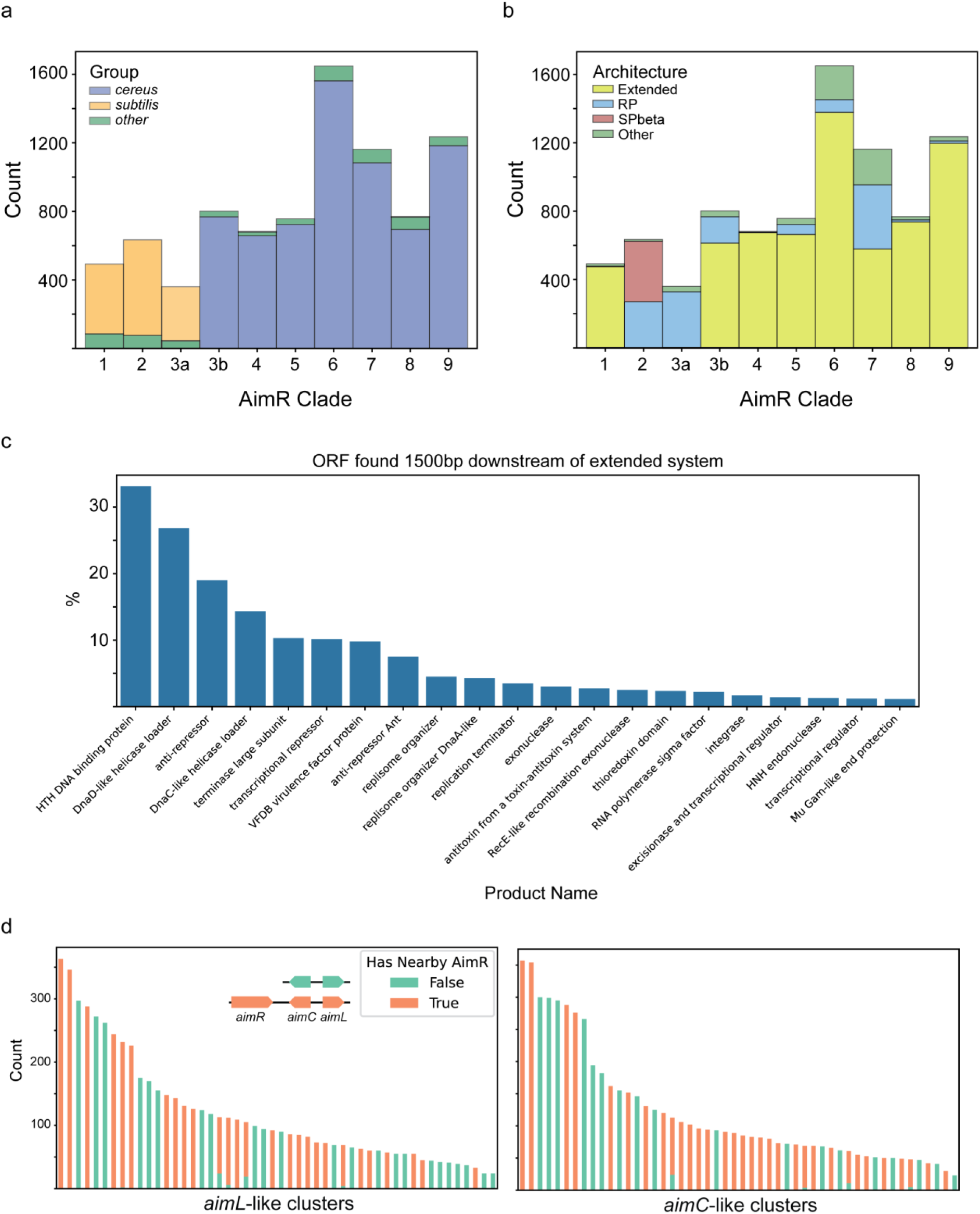
Additional data on arbitrium architectures and diversity. (a,b) Distribution of arbitrium coding species (a) and architectures (b) across arbitrium clades. (a) Shown are the association of different bacillus groups (*subtilis*, *cereus*, *other*), for different AimR clades, as defined in ref ^11^ with a single change – clade 3 was split into two phylogenetic sub-clades. This distinction split this clade into subtilis-dominated and cereus-dominated. (b) The different major architectures shown in Fig. 1b, but now for each clade. Clearly, SPbeta is found only in clade 2 (though the RP architecture is also found there). The extended architecture is a large majority in all clades except 2,3a. (c) putative co-cistronic genes, 1.5 kbp downstream of *aimL*, were annotated in all extended systems using PHANOTATE^57^. Shown are the frequency of appearance of the top categories. (d) homology clusters of *aimL*-like and *aimC*-like pairs and their positive (orange) or negative (green) association with arbitrium systems. HMM were made for AimC and AimL based on the extended systems in which they were identified. An independent search was then performed for pairs of *aimC*-like and *aimL*-like genes in the correct architectures (left, inset) across the same Bacillus genomes. Each identified pair was marked as associated with AimR (orange) or not (green). *aimC*-like and *aimL*-like gene products were then clustered using DIAMOND^29^ with 80% similarity. Shown are the 50 largest clusters. The unique color of almost all clusters suggest that *aimC*-like-*aimL*-like pairs associated with arbitrium are distinct from those not associated with it, suggesting that *aimC* and *aimL* are strongly linked with the extended arbitrium systems.

**Supplementary Figure 2:**
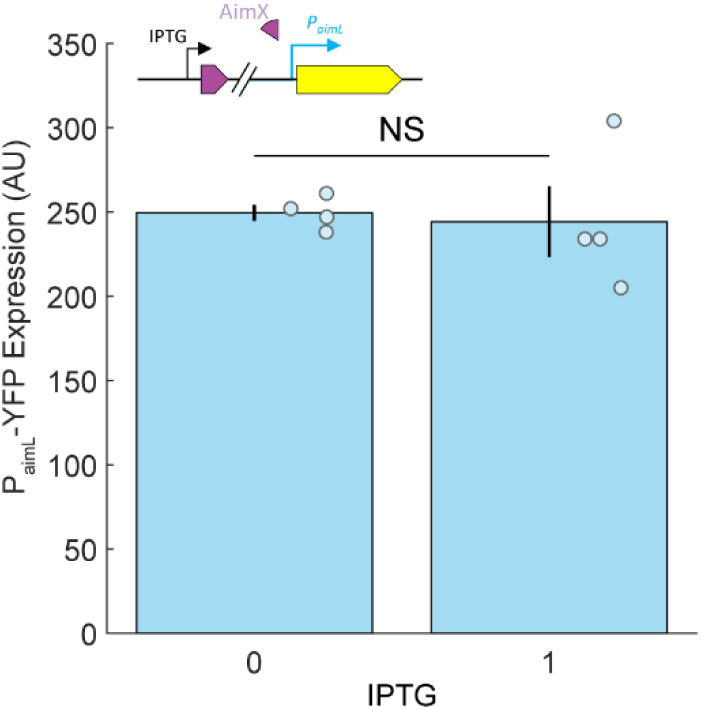
Expression of a P*_aimL_*-YFP reporter in a strain coding for an IPTG-inducible *aimX* but not for *aimC*. Shown are the reporter levels with and without IPTG. No significant difference in expression is found. Compare with Fig. 2d, where the same construct lead to drastic changes in expression in the presence of *aimC*.

**Supplementary Figure 3:**
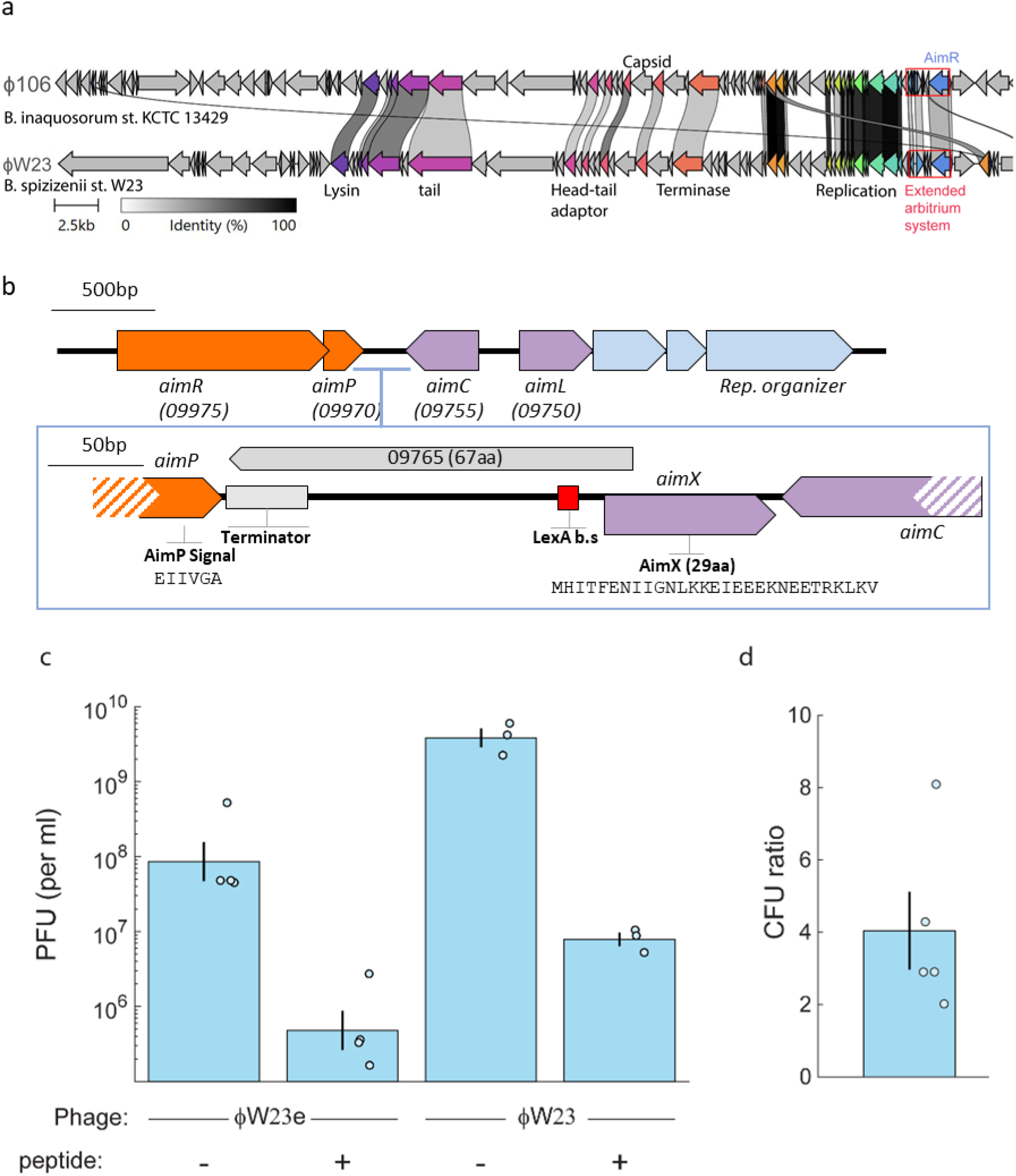
Characterization of phage ϕW23. (a) a comparative map of phages ϕ106 of *Bacillus inaquosorum* strain KCTC 13429 and ϕW23 from *Bacillus spizizenii* strain W23. Lines indicate homology according to the legend. Major homologous regions or genes are annotated. The extended arbitrium systems are marked by a red box. (b) a closeup on ϕW23 extended arbitrium system. Annotated are the different genes and their locus tag number on W23 genome (accession CP002183.1). Note the difference in AimX sequence from ϕ106, the lack of overlap with *aimC*, the different arbitrium peptide and the presence of an uncharacterized putative ORF of 67aa length between *aimC* and *aimP* in the direction of *aimC* (enlargement, gray color) (c) Phage titer count of the erythromycin resistance marked (ϕW23e) and unmarked (ϕW23) phages two hours after infection in the same conditions (OD 0.3, MOI 0.1) with, or without the ϕW23 arbitrium peptide. (d) CFU count of the ratio between the number of lysogens formed 30 minutes after infection with and without addition of the peptide. This ratio is significantly different than 1 (p=0.005, t-test of logarithm). Notably, the ratio after two hours of infection was non-significantly different than 1. (e) Specific mutations in the ϕW23;*aimX*^stop^ allele.

**Supplementary Figure 4:**
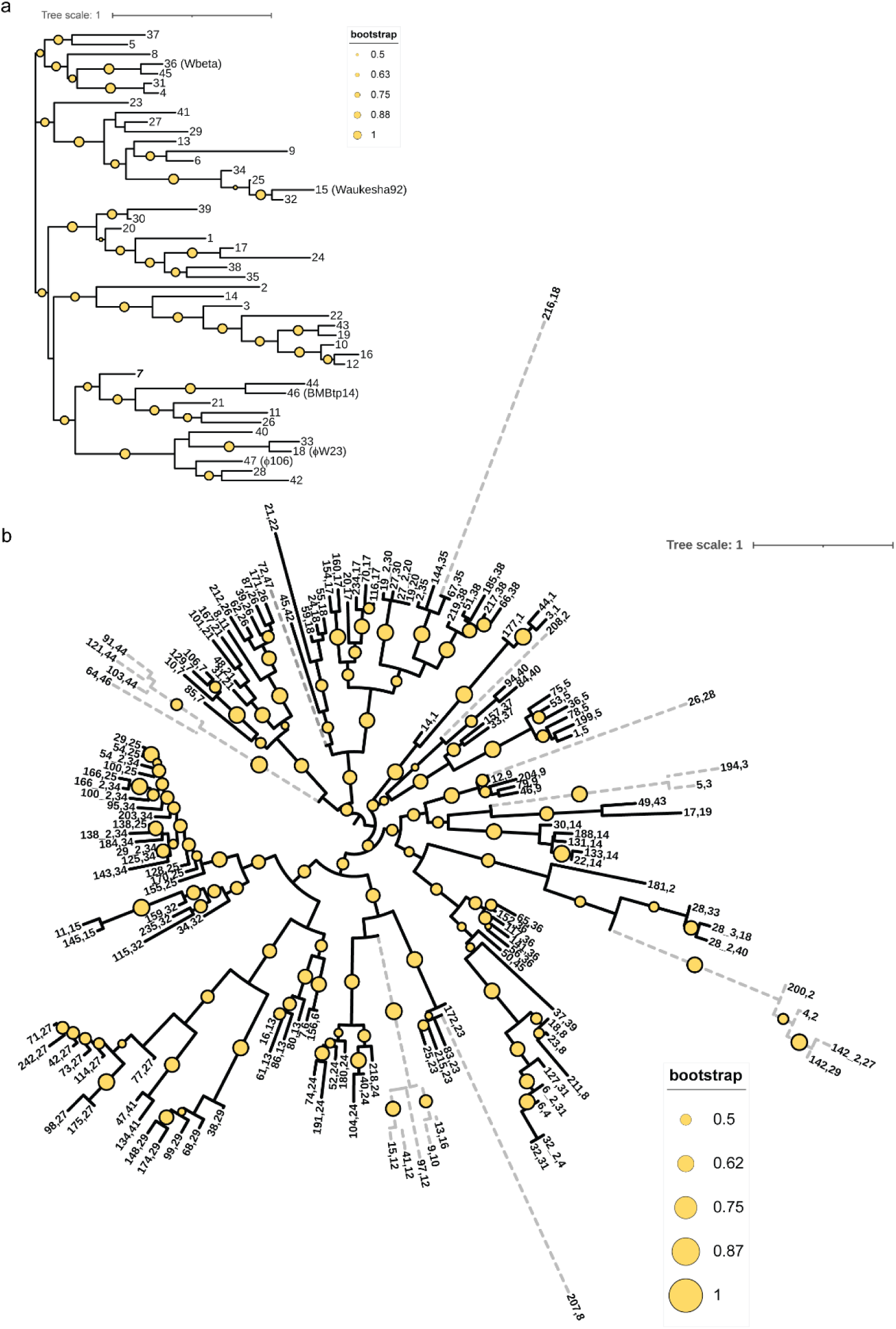
Phylogenetic trees of AimC, AimX and the relation between them. Shown is the phylogenetic tree of (a) AimC and (b) AimX, based on a single representative taken from each cluster of 80% similarity (as defined by diamond ^29^). The figure in a is the same as in Fig. 4a, but clusters numbers (ordered by number of elements in each cluster) are given, Each leaf in (b) is marked both by its own cluster number and the corresponding cluster number of AimC proteins which are found associated with the AimX in that cluster. Note that similar AimX clusters are associated with the same or similar clusters of AimC. Note the scale in (b). Branches whose length is >1 are dashed, pointing to a large uncertainty in their position. Bootstrap results are marked by circle sizes as in the legends in a,b.

**Supplementary Figure 5:**
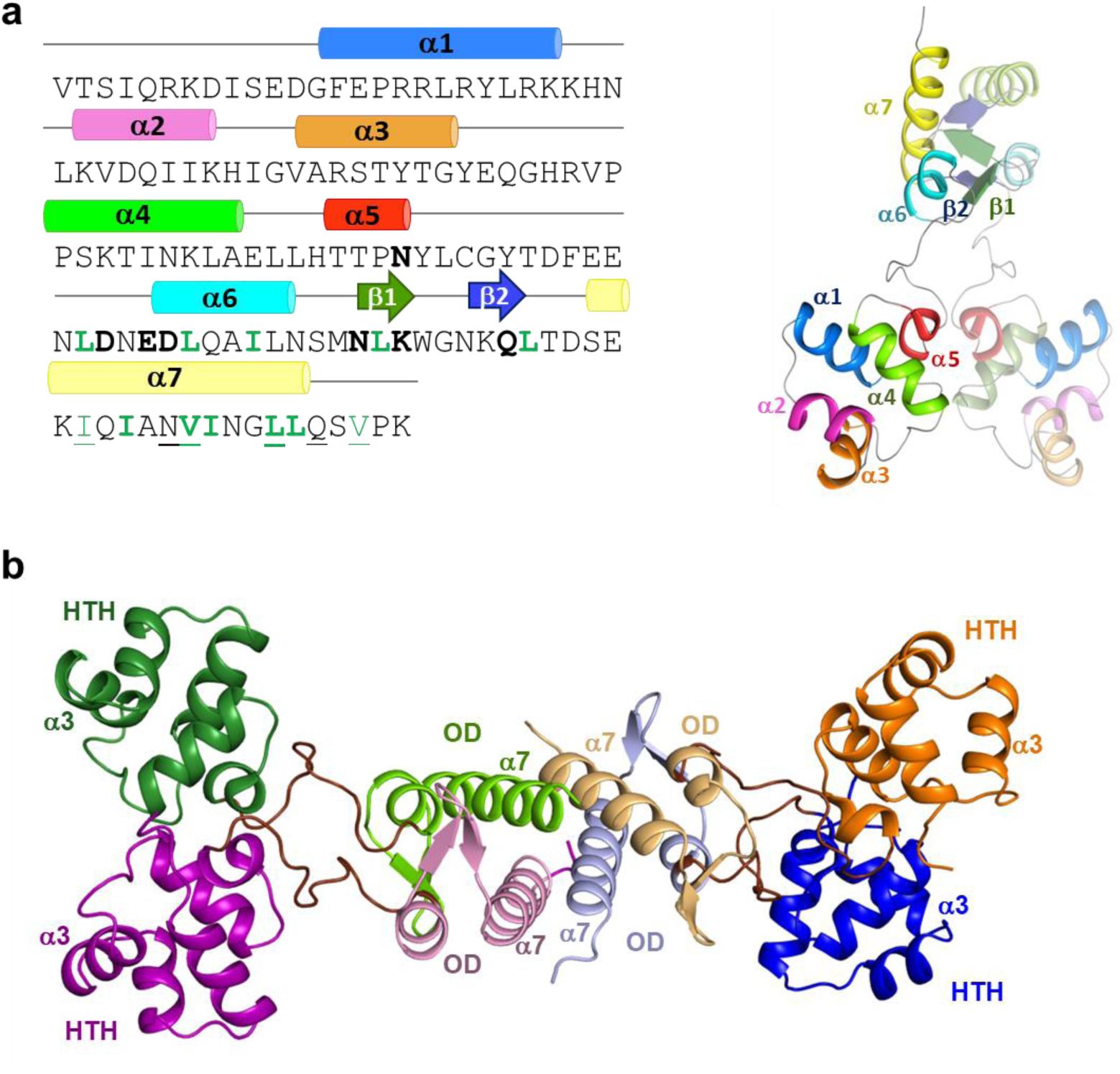
AimC_BMBtp14_ Structure and quaternary organization. A) Sequence of AimC_BMBtp14_ (left) and ribbon representation of the dimer structure (right) highlighting the secondary structure elements (above the sequence) in different colours. In the structure one of the monomers is drawn semi-transparent. Residues involved in dimerization are highlighted in bold, and residues involved in tetramer formation are underlined. Hydrophobic residues involved in interactions are coloured in green. To form the dimer the helices align generating a four-helix bundle closed laterally by the β-hairpins. This four-helix bundle is stabilized by hydrophobic interaction and hides ∼1100 Å^2^ surface by monomer. b) Ribbon representation of AimC_BMBtp14_ tetramer formed in the asymmetric crystal unit. Monomers are shown in purple, green, red and orange with darker colors for the HTH domain and lighter for the oligomerization domain (OD). Tetramerization is carried out exclusively by the reciprocal interactions of the four c-terminal α7 helices, providing each one a smaller surface of ∼390 Å^2^. DNA-binding α3 and tetramerization α7 helices are labelled on each of the monomers.

**Supplementary Figure 6:**
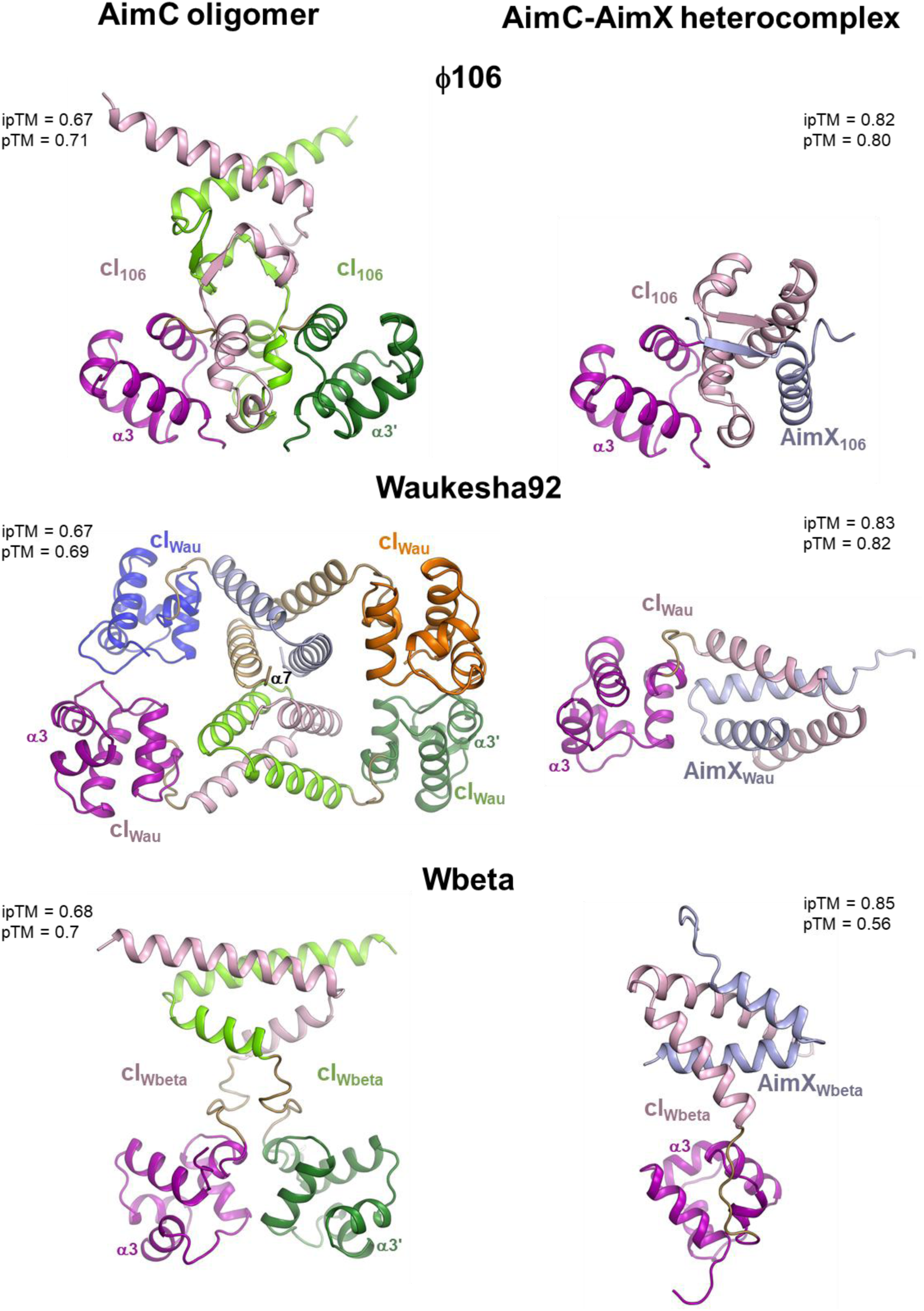
Alphafold 3^33^ models of AimC multimer and AimC-AimX dimer structure for additional AimC-AimX systems. Shown are the predicted structures predicted by alphafold 3 for phages ϕ106, Waukesha92 and Wbeta. Note that Waukesha92 AimC multimer is predicted to be a tetramer while the other two are predicted to form dimers. ipTM and pTM scores are given for each structure.

